# Host Induced Gene Silencing of the *Sclerotinia sclerotiorum ABHYDROLASE-3* gene reduces disease severity in *Brassica napus*

**DOI:** 10.1101/2021.11.26.470135

**Authors:** Nick Wytinck, Dylan J. Ziegler, Philip L. Walker, Daniel S. Sullivan, Kirsten T. Biggar, Deirdre Khan, Solihu K. Sakariyahu, Olivia Wilkins, Steve Whyard, Mark F. Belmonte

**Author notes:** Corresponding Author: Dr. Mark F Belmonte, Department of Biological Sciences, University of Manitoba, Winnipeg, Manitoba, R3T 2N2, Canada. 1-204-474-8556.

## Abstract

*Sclerotinia sclerotiorum* is a pathogenic fungus that infects hundreds of crop species, causing extensive yield loss every year. Chemical fungicides are used to control this phytopathogen, but with concerns about increasing resistance and impacts on non-target species, there is a need to develop alternative control measures. In the present study, we engineered *Brassica napus* to constitutively express a hairpin (hp)RNA molecule to silence *ABHYRDOLASE-3* in *S. sclerotiorum*. We demonstrate the potential for Host Induced Gene Silencing (HIGS) to protect *B. napus* from *S. sclerotiorum* using leaf, stem and whole plant infection assays. The interaction between the transgenic host plant and invading pathogen was further characterized at the molecular level using dual-RNA sequencing and at the anatomical level through microscopy to understand the processes and possible mechanisms leading to increased tolerance to this damaging necrotroph. We observed significant shifts in the expression of genes relating to plant defense as well as cellular differences in the form of structural barriers around the site of infection in the HIGS-protected plants. Our results provide proof-of-concept that HIGS is an effective means of limiting damage caused by *S. sclerotiorum* to the plant and demonstrates the utility of this biotechnology in the development of resistance against fungal pathogens.

## Introduction

Fungal pathogens represent a worldwide threat to food security and cost producers billions of dollars in lost yield [1]. Traditionally, broad-spectrum agrochemicals have been used to limit the effects of these pathogens, but due to growing concerns of the impacts of these products on the agroecological environment as well as increasing incidences of fungicide resistance, there is a need to find more effective and sustainable solutions [2–4]. Breeding for genetic resistance to fungal pathogens is one approach at reducing our dependence on traditional fungal control strategies, but for necrotrophic fungi, genetic resistance is difficult to achieve [5]. Sclerotinia stem rot, caused by the fungal pathogen *Sclerotinia sclerotiorum,* is a disease that affects over 600 hosts, including *Brassica napus* [6]. The complex lifecycle, which involves several years of dormancy within the soil, a broad host range and multifaceted infection of all phyllospheric components of the plant, provides numerous challenges for effective control [6, 7]. Since no truly resistant cultivars of *B. napus* have been identified against *S. sclerotiorum,* there is an immediate need to develop novel crop protection technologies to control this fungal pathogen [8–10].

*S. sclerotiorum* requires a senescing nutrient source in order to germinate ascospores. In *B. napus* infection, it is the flower petal that facilitates germination. The infection can spread to leaf tissue as the flowers fall off, where the fungus is able to penetrate and colonize the vasculature of the plant [11]. Once inside the vasculature, the fungus travels laterally into the stem, penetrating the interior, and degrading the pith tissue, and ultimately causing lodging [4]. Due to the highly aggressive nature and speed of this infection, innate plant defense mechanisms are generally insufficient, and the host plant becomes overwhelmed by the fungus [4]. During their coevolutionary arms race, plants have developed a basal immune system that first detects pathogen-specific cell surface components through pattern recognition receptors (inducing pattern triggered immunity) and secreted pathogen-derived effectors by nucleotide-binding leucine rich repeat proteins (inducing effector triggered immunity) [12]. Detection of the fungal pathogen elicits downstream cellular responses including changes in hormone signalling, transcriptional reprogramming, and defense gene activation [13, 14]. In *B. napus*, pattern triggered immunity induces the synthesis of reactive oxygen species, changes in ion fluxes across cellular membranes, and increases in calcium (Ca^2+^) concentrations within cellular spaces [11, 15–17]. Calcium ions act as messengers triggering several cellular responses including the synthesis of phytoalexins, pathogen-related (PR) proteins, and programmed cell death of neighbouring cells [18]. Endogenous hormone changes further induce cell wall reinforcement through polysaccharide deposition such as callose around phloem sieve pores, activation of downstream defense genes and accumulation of secondary metabolites [19–21]. *S. sclerotiorum* does not exhibit a gene-for-gene response during interactions with the host unlike other *B. napus* pathogens such as *Leptosphaeria maculans* or *Plasmodiophora brassicae*, and therefore despite advances made in the understanding of the *B. napus – S. sclerotiorum* interaction, few control strategies using genetic tools have proven successful.

RNA interference (RNAi)-based technologies have emerged as a promising alternative control strategy for a growing number of fungal phytopathogens [22–24]. Given the sequence specificity of RNAi, application of these technologies could limit off-target effects [25]. RNAi-based crop protection technologies can be designed in two ways. First, through a topical approach where the double stranded RNA is applied as a spray (spray induced gene silencing; SIGS). Second, through genetic modification of the host plant to produce hairpin dsRNA molecules targeting the pathogen (host induced gene silencing; HIGS) [24]. Both methods have proven effective at limiting infections and disease symptoms of various fungal pathogens. SIGS is considered to be a more versatile technology, as the molecules can be applied to any host plant, they can be synthesized and produced in large quantities, and the technology may be more readily accepted in countries where genetically modified technologies are banned [26, 27]. HIGS on the other hand, is considered a more durable technology that provides constitutive protection against the pathogen and limits the amount of fungicide applied to the crop [28]. HIGS can be tailored to provide a novel source of genetic control that specifically targets the virulence of the pathogen, unlike many traditional engineering technologies, which instead aim to enhance the defense response of the plant [29, 30].

HIGS was first demonstrated in 2010 by Tinoco *et al.* [31] and Nowara *et al.* [32]. Tinoco *et al.* [31] engineered tobacco to express a β-glucuronidase (GUS) hairpin to specifically silence GUS transcripts in a GUS-expressing strain of *Fusarium verticilloides* while Nowara *et al.* [32] developed powdery mildew (*Blumeria graminis*)-tolerant wheat and barley . HIGS has also proven effective in protecting plants against biotrophic fungi such as *Puccinia triticina* [33], *Fusarium oxysporum* [34], hemibiotrophs such as *F. graminearum* [35] and necrotrophs such as *Botrytis cinerea* [36]. The mechanism behind HIGS, called cross-kingdom RNAi has been observed in plant-fungal interactions, where native small RNA molecules from the host plant are produced to minimize the virulence of the pathogen, or conversely, the pathogen can produce RNAs that reduce the ability of the plant to activate a defense response [30]. For example, Weiberg *et al.* [37] observed that *Botrytis cinerea* produces small RNA effectors that target and suppress host cellular immune machinery, and this transfer of small RNAs to the host could be inhibited by knocking down fungal *DICER-LIKE 1/2*. Cross-kingdom RNAi has also been observed in the *S. sclerotiorum-B. napus* pathosystem, where the host genes *SOMATIC EMBRYOGENESIS RECEPTOR KINASE 2* (*SERK2*) and *SNAKIN-LIKE CYSTEIN RICH PROTEIN 2* (*SNAK2*) are specific targets of *S. sclerotiorum* small RNAs [38]. Interestingly, Niehl et al. [39] found that another SERK, *SOMATIC EMBRYOGENESIS RECEPTOR KINASE 1* signalling is activated in *Arabidopsis thaliana* upon dsRNA detection through a pattern triggered immunity-like response. Cai *et al.* [40] demonstrated that *A. thaliana* plants could, in contrast, protect themselves from *B. cinerea* by transferring small RNA molecules within extracellular vesicles to the fungus to silence virulence gene expression. Koch *et al.* [41] confirmed that small RNAs derived from HIGS-based transgenes are also shuttled to invading fungi in this manner within the *A. thaliana-Fusarium graminearum* pathosystem. Thus, HIGS can exploit cross-kingdom RNAi to complement the other defenses within the plant against the pathogen.

While effective HIGS solutions have been developed for many economically significant pathosystems [31–36], one system that remains unreported is the *B. napus-S. sclerotiorum* interaction. The dsRNA-mediated knockdown of a predicted aflatoxin biosynthesis gene and pathogenicity factor, SS1G_01703 (*ABHYDROLASE-3*) proved to be highly effective at conferring plant protection using SIGS [42]. Here, we hypothesize that this gene target would be a good candidate to develop a HIGS-based control system in *B. napus*. In the current study, we demonstrate that *B. napus* expressing hairpin (hp) RNAs targeting the *S. sclerotiorum ABHYDROLASE-3* gene (SS1G_01703) produce transgene-derived small RNAs and reduced the transcript levels of SS1G_01703. Restriction of *S. sclerotiorum* infection through knockdown of *ABHYDROLASE-3* together with global changes in plant defense resulted in improved plant health and seed set. Taken together, HIGS provides a genetic tool that works with the innate defense response of the host plant to reduce *S. sclerotiorum* infection in *B. napus* through a targeted RNAi approach.

## Results

### Confirmation of transgene expression

*B. napus* cv. Westar was transformed to express a hpRNA targeting *S. sclerotiorum ABHYDROLASE-3* (SS1G_01703) under the control of a CaMV 35S promotor (Bn35S::SS1G_01703RNAi hereby referred to as BN1703) (Fig. 1a, Fig. S1). *S. sclerotiorum ABHYDROLASE-3* is predicted to be involved in aflatoxin synthesis due to amino acid similarity to aflatoxin biosynthesis genes in related fungi (Table S3). Five different independent insertion lines were propagated for three generations. The transgene copy number in these lineages, determined by qPCR of the transgene relative to the single copy gene, High Mobility Group protein *HMG-I/HMG-Y* (BNHMG I/Y, *BnaA06g34780D*) [43], ranged between two and three copies in each genome (Fig. 1b). We also examined the expression levels of the gene encoding the endonuclease *DICER-LIKE 2,* which has been shown to process hpRNAs derived from transgenes [44]. qRT-PCR of *B. napus DICER-LIKE 2* (*BnaA05g32540D*) was 150 to 350 times more abundant in the transgenic lines compared to their untransformed counterparts (Fig. 1c). BN1703.2 demonstrated the highest abundance of DCL2A and transgene copies. To confirm that the hpRNAs were processed into small RNAs in BN1703.2, we sequenced small RNA populations genome-wide in uninfected and infected stems. Small RNAs specific to the SS1G_01703 transgene were indeed present prior to and during infection; none of these small RNAs were detected in untransformed plants. Within the pool of small RNAs aligning to the transgene, sequence reads aligning to specific regions of the transgene were found to be in greater abundance than others (Fig. S2). Using linear-hairpin variable primer qRT PCR (Fig. 1d), the levels of one of these abundant siRNAs was quantified, and observed to be more abundant in the BN1703.2 lines, relative to the other lineages (two times more abundant than in BN1703.1), and not detectable in the untransformed control (Fig. 1e). qRT-PCR analyses of SS1G_01703 transcript levels in both untransformed and BN1703.2 stem tissues revealed a strong and persistent reduction of SS1G_01703 transcripts in the BN1703.2 plants over the course of 7-days infection, compared to the untransformed plants (Fig. 1f). Thus, the successful expression of a hpRNAi transgene construct resulted in gene silencing of the *S. sclerotiorum ABHYDROLASE-3* transcript.

**Fig. 1.**
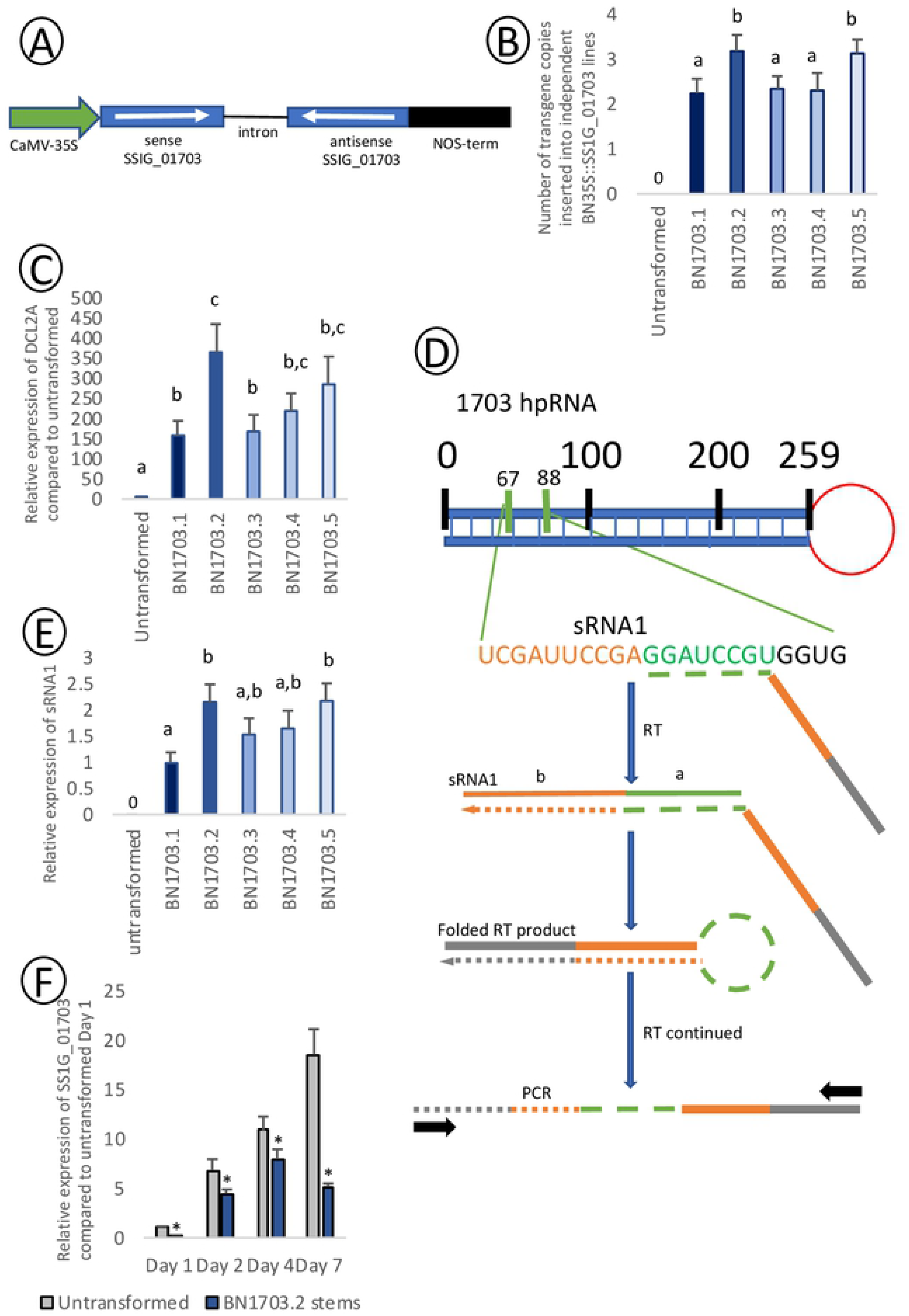
BN1703 encodes, expresses and processes hpRNA targeting SS1G_01703. (A) Schematic map of the BN1703 RNAi cassettes within the pHELLSGATE8 vector. The intron-hairpin interfering cassette is under the control of the 35S promotor of Cauliflower mosaic virus. The 259 bp fragment of the *ABHYDROLASE-3* gene coding sequence from *S. sclerotiorum* (SS1G_01703) was directionally cloned to generate sense and antisense arms flanking an intronic spacer. NOS (nopaline synthase) provides selection against kanamycin. (B) Estimated number of BN1703 copies inserted into *B. napus* transformants relative to single copy gene BNHMG I/Y. (C) Relative expression of BNDCL2A in *B. napus* transformants relative to untransformed plants and normalized to the housekeeping reference BNATGP4. (D) Schematic diagram of the amplification of sRNA1 from 1703 hpRNA using linear variable hairpin qPCR through the reverse transcribed extension of a hairpin primer. (E) Transcript abundance of sRNA1 using linear variable hairpin qPCR of *B. napus* transformants. Data for nucleic acid quantifications represents three biological replicates with error bars representing standard error. Statistical differences calculated using one way ANOVA (with significance of p<0.05). (F) Relative expression of SS1G_01703 in infected untransformed and BN1703.2 *B. napus* stems 1, 2, 4, and 7 days post-inoculation demonstrating transcript knockdown within BN1703.2 stems. Asterisks represent statistical differences from the untransformed control (one-tailed t-test with Bonferroni correction, p<0.05 from three biological replicates).

### HpRNA production in *B. napus* host targeting SS1G_01703 significantly reduces *S. sclerotiorum* infection

Next, we challenged BN1703 lines with *S. sclerotiorum* using a petal inoculation method on leaf tissues [11] and the mycelial plug method on stems [45] (Fig. 2a). Within the T2 and T3 generations, reductions in lesion sizes were observed in both stem and leaf infection assays (Fig. S3a). In addition to highest copy number (Fig. 1b), DCL2A (Fig. 1c) and sRNA1 abundance (Fig. 1d), BN1703.2 also showed the greatest reduction in fungal lesion size (Fig. 2b). Line BN1703.2 also accumulated significantly less fungal rDNA in the stem over the course of infection compared to the untransformed control (Fig. 2c). Lactophenol ‘cotton’ blue staining, which detects fungal chitin [46], was used to examine plant stem infections of *S. sclerotiorum*. Transverse sections of untransformed plant stems demonstrated that the fungus successfully colonized the interior pith region 7 days post inoculation. Lactophenol staining within the secondary xylem indicated radial infiltration of the fungus along medullary rays in the untransformed plants (Fig. 2d). Conversely, the *S. sclerotiorum* infiltration within the BN1703.2 plants does not proceed into the secondary xylem at 7dpi as observed in untransformed plants, and instead is observed no further inward than the cortex and secondary phloem. (Fig. 2e).

**Fig. 2.**
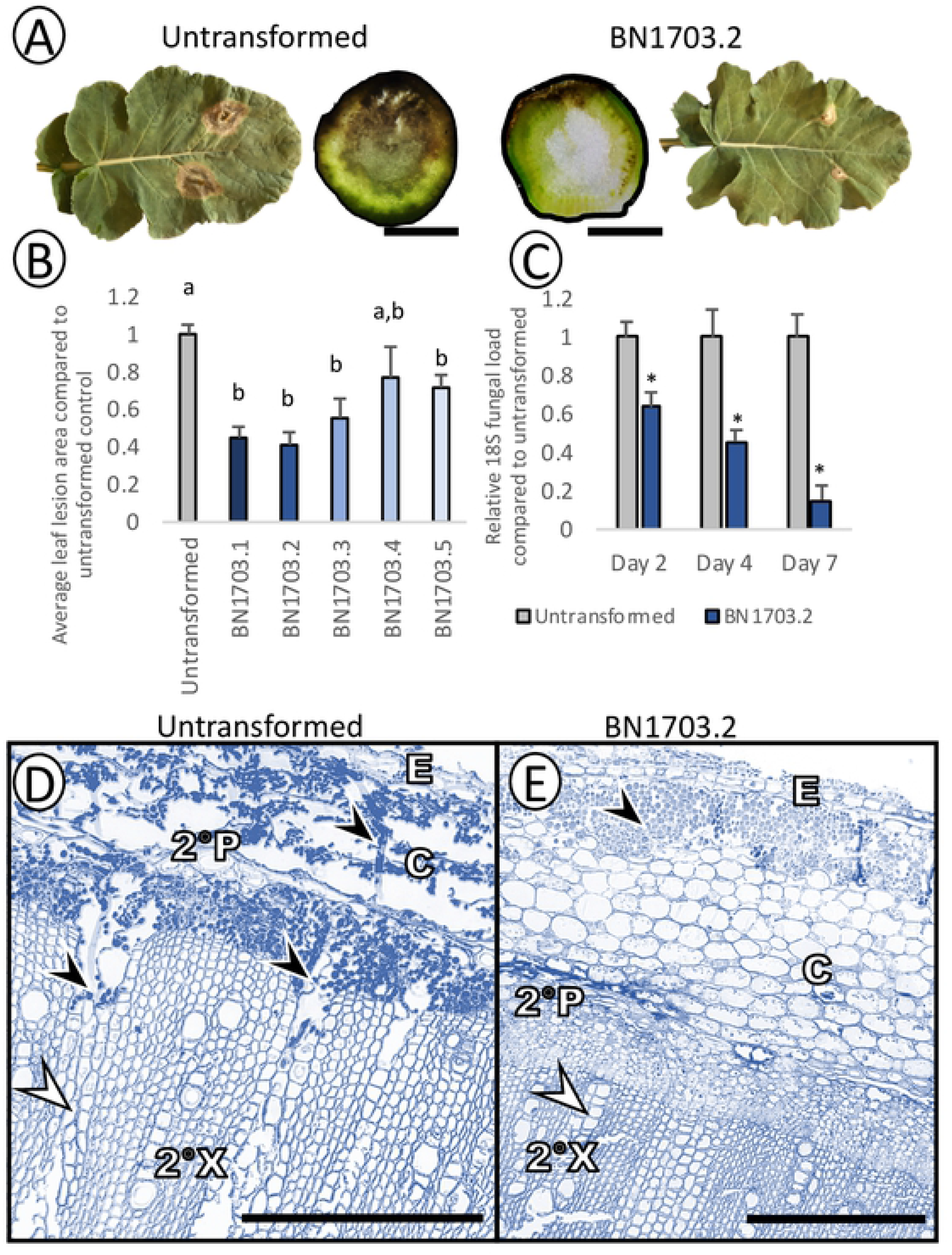
BN1703.2 confers significant protection against *S. sclerotiorum* infection. (A) Representative images of infected untransformed and BN1703.2 *B. napus* leaves and stem cross-sections 3 and 7 days post-inoculation respectively. Scale bars represent 1 cm for stem cross sections. (B) Relative leaf lesion area of infected *B. napus* transformants compared to untransformed leaf lesions. Data represents at least ten leaf lesions per line with error bars representing standard error. Statistical differences were tested with a one-way ANOVA (with significance of p<0.05), where significant differences are denoted with differing letters. (C) Relative fungal load of infected stem lesions 2, 4, and 7 days post inoculation. 18S rDNA abundance was quantified for untransformed and T2 BN1703.2 stem lesions. Asterisks represent statistical differences from the untransformed control (one-tailed t-test with Bonferroni correction, p<0.05 from 3 biological replicates). Lactophenol blue staining of stem transverse-sections 7 days post-inoculation of untransformed (D) and BN1703.2 (E) stems.

Untransformed stems show extensive colonization and degradation of the epidermis (E), cortex (C) and secondary phloem (2°P) and penetration into the secondary xylem (2°X) through medullary rays. BN1703.2 stems show colonization of the epidermis (E) and cortex (C), although they remain intact. Secondary phloem (2°P) and xylem (2°X) remain uncolonized. Black arrowheads indicate *S. sclerotiorum* hyphae highlighted by lactophenol blue stain and white arrowheads indicate medullary rays. Scale bars represent 250 μm and micrographs are representative images of at least three biological replicate stems.

### Host induced gene silencing of SS1G_01703 confers tolerance to *S. sclerotiorum* across the plant lifecycle

Quantification of the SS1G_01703-specific siRNAs confirmed that the transgene was expressed and processed into the hairpin siRNA in flowers, leaves and stems in uninfected BN1703.2 plants, the three prominent locations of *S. sclerotiorum* infection on *B. napus* (Fig. 3a). *B. napus* plants at the 30% flowering stage were sprayed with a concentrated (1x10^6^ spores/mL) solution of *S. sclerotiorum* ascospores, coating the whole plant, and disease progression was surveyed over 7 days under 90-100% humidity. After 7 days, untransformed plants were more affected by *S. sclerotiorum*. Petal forming lesions and total sclerotia were reduced by 70 and 60 percent respectively in infected BN1703.2 plants compared to the untransformed counterparts. Finally, total seed mass of the plants after maturation increased three-fold in line BN1703.2 compared to that in untransformed (Fig. 3b). In untransformed plants, most leaves became necrotic and the stem was weakened by fungal infection at several sites (Fig. 3c). In contrast, while lesions were present on leaves and stems of line BN1703.2, these plants showed less necrosis and were able to continue through to maturation (Fig. 3c).

**Fig. 3.**
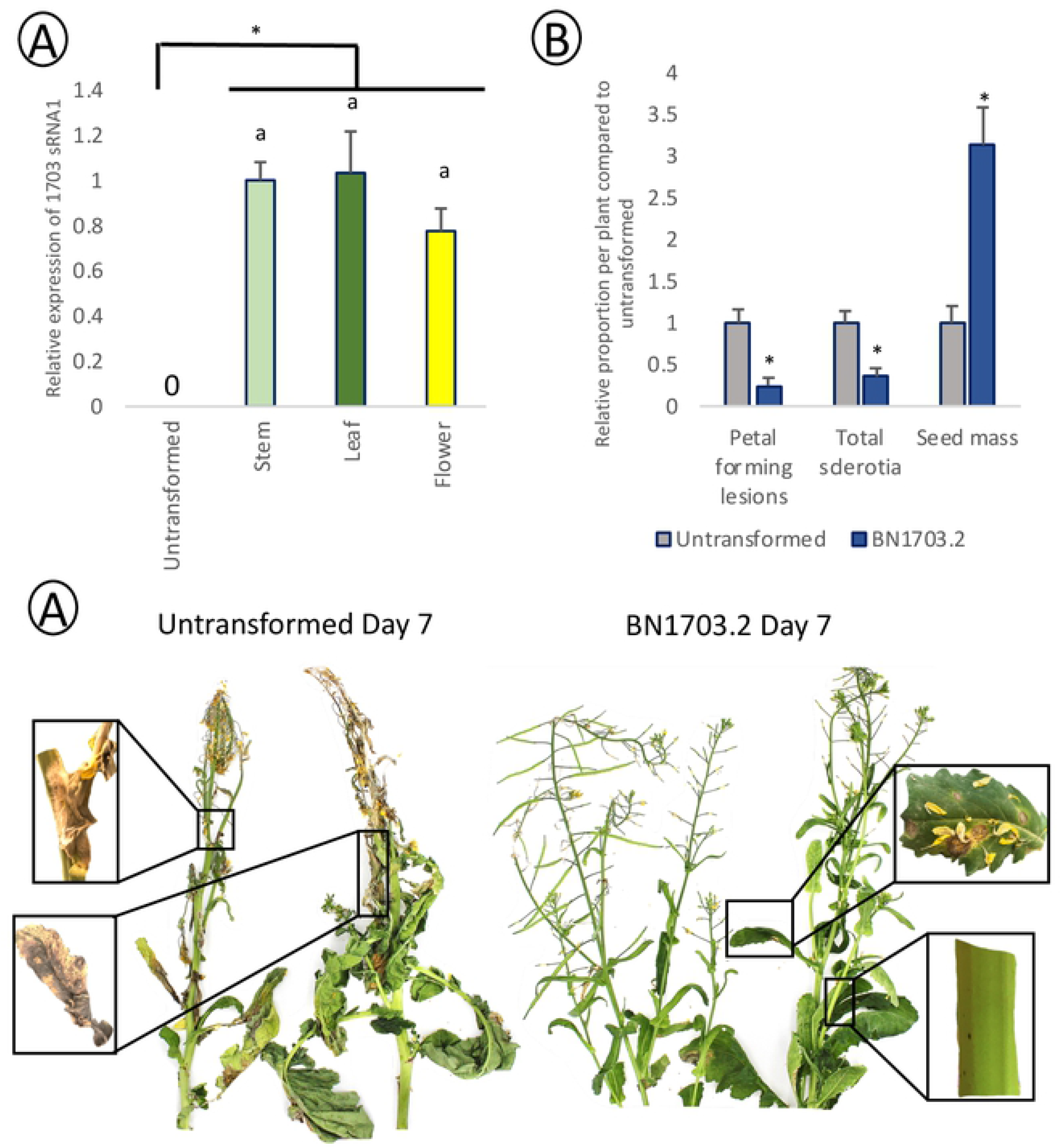
BN1703.2 confers whole protection up to plant maturity. (A) Relative abundance of sRNA1 in BN1703.2 uninfected stems, leaves and flowers to demonstrate transgene expression in these organs. Statistical differences were tested with a one-way ANOVA (with significance of p<0.05), where significant differences are denoted with differing letters. (B) Total petal forming lesions per plant were quantified after 7 days of incubation in a humidity chamber after spraying. Plants were then allowed to mature fully before total sclerotia and seed mass were quantified per plant. Data represents at least ten plants per cultivar and asterisks represent statistical differences from the untransformed control (one-tailed t-test with Bonferroni correction, p<0.05). (C) Representative images of ascospore spray infected untransformed and transgenic BN1703.2 plants after seven days of incubation in a humidity chamber after spraying. Untransformed plants show extensive infection on flowers and leaves and several sites of lesion progression into the stem. BN1703.2 plants show infection of flowers and leaves however, minimal lesion progression upon the leaves and stem.

### RNA sequencing of HIGS-based plant protection

To better understand how both transformed and untransformed *B. napus* respond to *S. sclerotiorum*, we profiled global gene activity in the stem at the plant-pathogen interface (Fig. 4a, Data S1). RNA sequence reads were mapped to the reference genomes of *B. napus* cv. Westar [47] and *S. sclerotiorum* [48] and differentially expressed genes (DEGs) compared to day 0 of the uninfected stems. We identified 37778 *B. napus* DEGs throughout the experiment (21405 upregulated and 16373 downregulated). The most substantial transcriptional response in terms of differential expression occurred at 1 dpi in BN1703.2 (6654 upregulated and 5692 downregulated) and the most minor transcriptional response occurred at 7 dpi also within BN1703.2 (3787 upregulated and 2388 downregulated). In total, we identified 1594 shared DEGs upregulated within all four comparisons (darkest shaded bar), indicating a common, coordinated response regardless of host genotype or stage of infection.

**Fig. 4.**
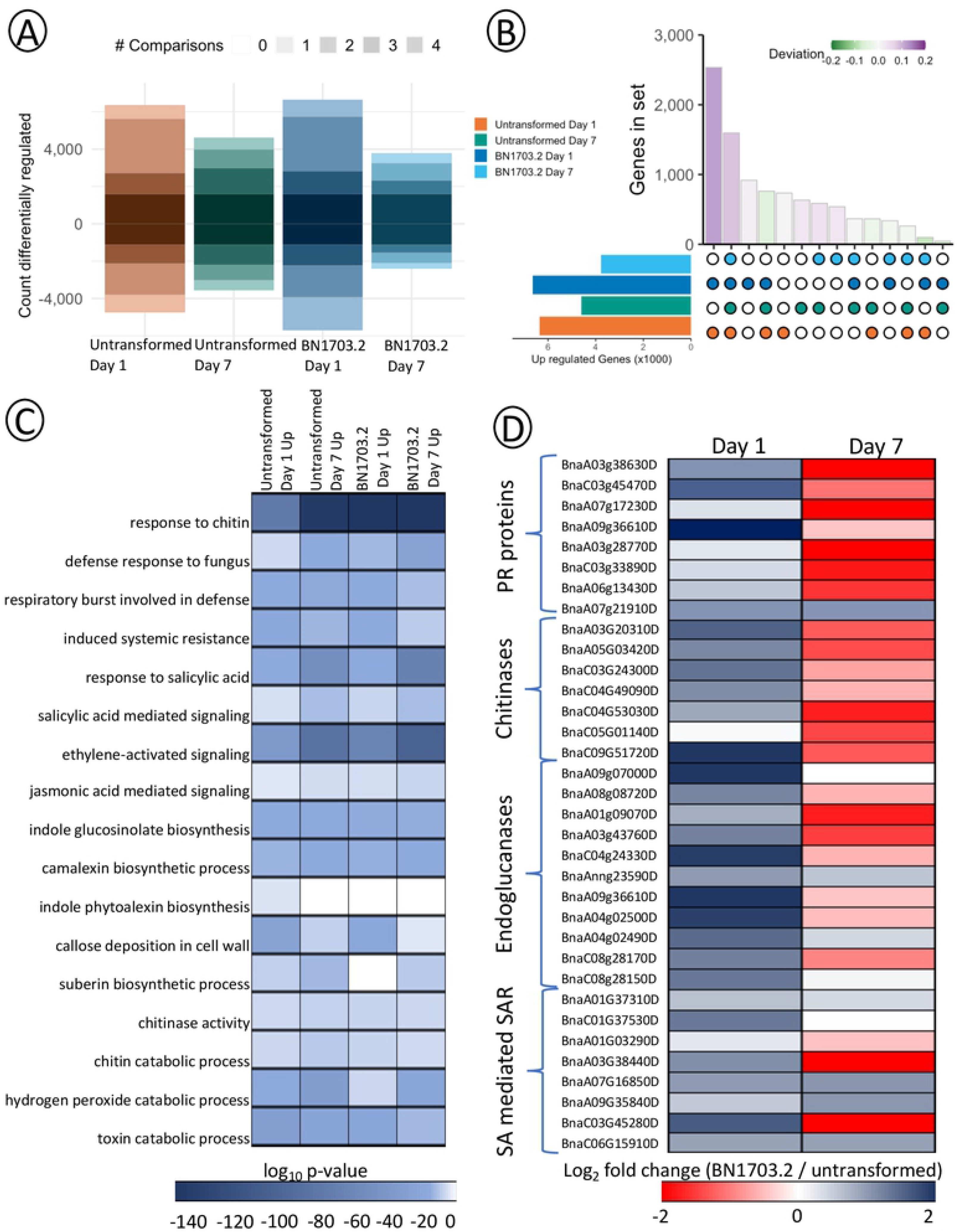
Transcriptomic characterization of plant defense to *S. sclerotiorum* in BN1703.2. (A) Stacked bar plot displaying the number of *B. napus* genes differentially expressed in untransformed and transgenic BN1703.2 infected stems 1 and 7 days post-inoculation and compared to the uninfected control (false discovery rate<0.01). Darker shades of color represent higher numbers of patterns shared between treatments. (B) Upset diagram showing the sizes of gene sets induced by each combination of patterns. The bars for the set sizes are coloured by the deviation from the size predicted by random mixing. (C) Heatmap of significantly enriched GO terms of untransformed and BN1703.2 days 1 and 7 compared to uninfected from total DEG. GO terms are considered statistically significant if the hypergeometric P-value<0.001. (D) Heatmap of a subset of genes encompassing significantly enriched GO terms within BN1703.2 during infection. GO terms are considered statistically significant if the hypergeometric P-value<0.001.

Principal component analysis of detected transcripts in *B. napus* grouped samples based on the timing of infection and genotype suggesting both genotypes underwent significant shifts in transcript abundance in response to infection (Fig. S4a). *S. sclerotiorum* reads clustered based on timing of infection (Fig. S4b). To determine whether gene sets were induced specifically by either genotype or time of infection, we identified *B. napus* DEGs induced by all potential combinations of patterns (Fig. 4b). This analysis demonstrated that coordinated DEG sets upregulated between both genotypes at 1 dpi and common DEG shared between both genotypes at both 1 dpi and 7 dpi were most abundant of all tested combinations. However, specific DEGs to BN1703.2 at 1 dpi were the next most abundant, and therefore encompassed the most unique DEGs of any individual comparison. Upregulated DEGs shared between untransformed and BN1703.2 at 7 dpi were the least abundant of all possible combinations, thus indicating that the responses of these genotypes are vastly different later in infection. Taken together, BN1703.2 had the strongest transcriptional response to *S. sclerotiorum* at 1 dpi.

Next, to investigate biological processes, molecular functions, and cellular components contributing to the *B. napus* defense response, we performed Gene Ontology (GO) term enrichment on all upregulated differentially expressed gene sets [49] (Fig. 4c). GO terms were considered significantly enriched if the hypergeometric P-value was <0.001. We aimed to uncover GO terms that were uniquely enriched within each genotype and timepoint in addition to those terms that were shared between all timepoints and genotypes. For example, biological processes such as toxin catabolism, systemic acquired resistance through salicylic and jasmonic acid signalling, production of glucosinolates and phytoalexins, callose and suberin deposition in addition to fungal cell wall degradation were all enriched within the BN1703.2 and untransformed plant gene sets relative to uninfected plants (Fig. 4c). We then compared the transcript abundance values for each of the individual genes encompassed by these biological processes GO terms between untransformed and BN1703.2. Interestingly, while many of these genes are differentially expressed during both untransformed and BN1703.2 infections, we see large quantitative differences in gene activity, especially at 1 dpi (Fig. 4d). In BN1703.2, there was greater transcript accumulation of pathogenesis related proteins (*PR1-PR5)*, 1.5-4-fold increase in abundance; PR proteins are induced as part of the systemic acquired resistance response. Additionally, a suite of chitinases and endogluconases responsible for fungal cell wall degradation, in particular *CHB4* and *BETA 1,3 ENDOGLUCONASE,* were overrepresented within the transgenic plants (2.5-4.5-fold increase). SA and SA-mediated signalling has been shown to play a significant role in early defense responses against *S. sclerotiorum* in *B. napus* [50]. Genes associated with the generation of SA-activated defense molecules such as camalexin and scopoletin (*FERULOYL COA ORTHO-HYDROXYLASE*, *WRKY TRANSCRIPTION FACTOR 70*), two excreted secondary metabolites with antifungal properties, also showed increased activity in BN1703.2 plants at 1 dpi (1.5-2.5-fold increase). By 7 dpi, this trend had reversed, with untransformed plants showing higher levels of defense gene transcripts as the infection progressed. The pathogenesis-related protein, *PR1*, (*BnaC03g45470D*), is another common marker for salicylic acid signalling. The increases in transcript levels of *PR1* in the RNA-seq dataset reflect closely to the qRT-PCR analyses which showed a 1.5 to 2-fold increase in transcript accumulation of this gene 1-4 dpi in BN1703.2 plants compared to untransformed plants (Fig. S3b). In uninfected stems, we observed no difference in *PR1* expression between BN1703.2 and untransformed plants through qRT-PCR providing evidence for no premature transcriptional activation of plant defense responses to the dsRNA transgene insertion (Fig. S3b).

Next, we examined global gene activity in *S. sclerotiorum* on both transformed and untransformed *B. napus* (Fig. S5). At day 1 post infection, significantly fewer transcripts encoding plant cell wall degrading enzymes, including cellulases, endo-glucanases, exo- glucanases and beta-glucanases were 50-80% less abundant while hemi-cellulases such as endo-beta-xylanases, beta-mannosidases, alpha-xylosidases, alpha-galactosidases and alpha-I- arabinofuranosidases wer 90% less abundant in BN1703.2 compared to their untransformed counterparts. The expression of specific proteases such as *CALPAIN FAMILY CYSTEINE PROTEASE* and *SUBTILISIN-LIKE SERINE PROTEASE,* involved in the degradation of pathogenesis related proteins and the suppression of host plant immunity, were 2-3 fold higher in BN1703.2 at 1 dpi compared to the same point in the untransformed control stems of *B. napus*. Genes involved in detoxification, particularly glutathione S-transferases, were also enriched 1.5-2-fold within transgenic plants compared to untransformed plants at 1 dpi. We observe a similar trend at 7 dpi with the gene activity of *S. sclerotiorum* within BN1703.2 infections as we did with the gene activity of *B. napus*. As the infection progresses, there is a reduction in the quantitative differences observed between BN1703.2 and untransformed infections.

### Activation of structural defense pathways in BN1703.2 in response to *S. sclerotiorum* infection

Transcripts of each of the ten *B. napus* genes encoding suberin biosynthesis enzymes were in low abundance in BN1703.2 plants at 1 dpi and increased between 2-7-fold higher than their untransformed counterparts by 7 dpi (Fig. 5a, Table S4). Suberin is an extracellular polymer component of the so-called vascular coating within stems and roots, providing strength, structural support and a possible barrier to pathogen entry (Fig. 5b). Scanning electron microscopy of infected BN1703.2 plant stems revealed a band running parallel to the length of the infection site on the stem surface within the secondary xylem (Fig. 5d). In BN1703.2 plants, the fungus did not appear to cross this band, whereas in untransformed plants, the fungus had heavily colonized the pith of the stems (Fig. 5c). Interestingly, we were also able to visualize a similar structure within infected stem cross-sections of the semi-resistant cultivar, *B. napus* cv. ZhongYou 821 (Fig. S6a). To gain insight into the composition of this vascular coating, we used a lipophilic stain (Sudan IV) to detect the presence of lipids such as suberin within infected stems of BN1703.2 and untransformed plants. In untransformed transverse hand-sections, there is sparse accumulation of the stain within the secondary xylem (Fig. 5e). In contrast, within BN1703.2 stems, we detected heavy deposits of Sudan IV stain localized within the xylem in a similar location to the electron dense band pattern observed using SEM (Fig. 5f). Staining transverse sections of BN1703.2 plants with Periodic Acid-Shiffs and toluidine blue did not, however, highlight the electron-dense ring observed in the SEM sections, which suggests that the ring observed is not composed predominantly of polysaccharides or lignins (Fig. S6b).

**Fig. 5.**
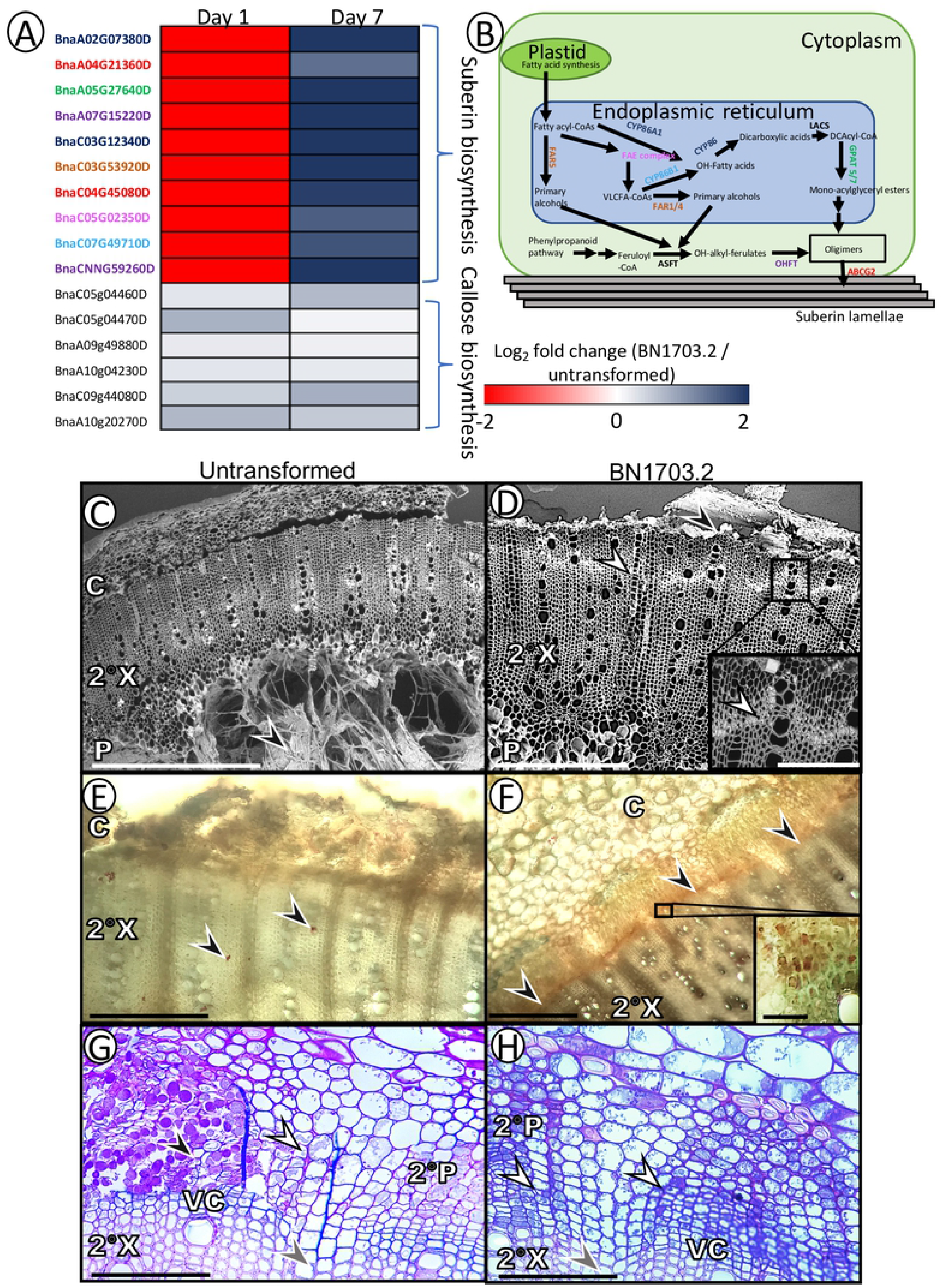
Structural differences between BN1703.2 and untransformed lead to enhanced protection from *S. sclerotiorum*. (A) Heatmap of genes involved in suberin and callose biosynthesis, which were enriched GO terms for BN1703.2. GO terms are considered statistically significant if the hypergeometric P-value<0.001. (B) A diagram of the biosynthesis pathway of suberin in plant cells. The enzyme font color corresponds to the *B. napus* gene ID in the heatmap in (A). Scanning electron micrographs of infected *B. napus* stems 7 days post-inoculation in untransformed (C) and BN1703.2 (D) genotypes. The untransformed stem shows extensive colonization throughout, particularly the pith (P) region. The BN1703.2 stem shows no colonization of the pith and a vascular coating within the secondary xylem (2°X). Scale bars represent 1 mm for untransformed and 0.5 mm and 50 μm for BN1703.2. Black arrowheads indicate *Sclerotinia* hyphae and white arrowheads indicate the vascular coating. Hand cut stem transverse sections stained with Sudan IV of untransformed (E) and BN1703.2 (F) stems 7 days post-inoculation. Red staining indicates deposition of suberin. The untransformed stem shows sparse Sudan staining while BN1703.2 has extensive deposition within the 2°X. Black arrowheads indicate Sudan IV stain and scale bars represent 150 and 50 μm for the lower and higher magnification respectively. Micrographs of stem transverse sections 7 days post-inoculation of untransformed (G) and BN1703.2 (H) stems stained with aniline blue. Aniline blue stains callose depositions dark blue-purple with the Periodic Acid-Schiff’s reagent stain staining plant and fungal cell walls bright pink. Untransformed stems show low levels of aniline blue stain in the secondary phloem (2°P) while BN1703.2 stems show heavy staining. Black arrowheads indicate *Sclerotinia* hyphae, white indicate callose, empty arrowheads indicate the vascular cambium and grey indicate the medullary ray. Scale bars represent 50 μm. All micrographs are representative of at least three biological replicates showing similar phenotypes.

Six genes associated with callose production (*CALLOSE SYNTHASES 3* and *7)* showed a 1.5-2-fold increase in transcript accumulation in BN1703.2 (Fig. 5a). Aniline blue was used to detect callose deposition within the secondary phloem. Within the BN1703.2 line, there appears to be abundant deposition of callose within the secondary phloem, particularly around the medullary ray regions proximal to the vascular cambium (Fig. 5h). Untransformed stems did not show the same pattern of deposition (Fig. 5g). Further, lumens of the sieve tube members proximal to the vascular cambium accumulate callose in BN1703.2 stems, which is not seen in the untransformed stems. Suberin and callose deposition appears to be a specific response to infection, as uninfected plants, whether transgenic or untransformed, showed no differences in the accumulation of these components (Fig. S6c-h).

## Discussion

Fungal control using RNAi-based strategies, and in particular, those using HIGS, provides a genetic tool to control fungal disease in crop plants (24,27). Here, we report HIGS control for *S. sclerotiorum* in *B. napus.* The *ABHYDROLASE-3* gene (SS1G_01703) was selected as a target for HIGS based on the previous work of McLoughlin *et al.* [42], which showed no orthologous gene targets in *B. napus* and knockdown of this target through foliar applications reduced *S. sclerotiorum* disease severity. While foliar applications of dsRNAs have shown promise in this pathosystem, a HIGS approach was hypothesized to confer more durable protection across the lifecycle of *B. napus*.

For HIGS to be effective, cross-kingdom trafficking must occur early in the infection, before it becomes systemic and overwhelms the plant [51, 52]. Ghag *et al.* [34] postulate that host-derived mobile signals such as small RNAs can gain entry into the fungal cytosol and induce gene silencing as cellular components between the host and pathogen are exchanged during the preceding biotrophic phase. The multistep lifecycle of *S. sclerotiorum* on *B. napus* provides opportunities for transgene-derived siRNA to be absorbed prior to necrotrophy. Wang *et al.* [22] and Wytinck *et al.* [53] have shown that fungal spores and emerging germ tubes in *B. cinerea* and *S. sclerotiorum* respectively, readily take in dsRNAs and siRNAs. Therefore, transgene-derived siRNAs may enter the fungal cell during the earliest stages of its lifecycle *in planta*. This early knockdown may lead to enhanced protection as the fungus transitions to necrotrophy and the infection cycle progresses. While cross-kingdom transfers of processed small RNAs between host plants and fungal pathogens have been described [40, 41], there exists the possibility that unprocessed long hpRNAs are being taken up by the fungus in a yet to be described mechanism. In transgenic maize expressing DvSnf7, sufficient quantities of un- diced, long dsRNA were sufficient to control the western corn rootworm, *Diabrotica virgifera* [54]. In that study, only long dsRNAs greater than 60 base pairs were shown to enter the gut cells of the insects to induce RNAi, while 21 bp siRNA molecules did not show knockdown of the target mRNA. In fungi, multiple studies have shown that both long dsRNAs and siRNAs can induce target mRNA knockdown. For example, in *B. cinerea,* both exogenous *DCL1, DCL2* dsRNA and siRNA caused similar levels of transcript knockdown *in vitro*, while siRNAs designed against *CHITIN SYNTHASE* in *Macrophomina phaseolina* were effective at suppressing fungal growth and inducing significant gene knockdown *in vitro* [22, 55]. However, while reduction in accumulation of target gene transcript provides an indication of RNAi-based silencing, examination of the degradome would be required to provide direct evidence for target cleavage.

While our initial hypothesis was that the constitutive knockdown of *ABHYDROLASE-3* would be sufficient to cause a reduction in *S. sclerotiorum* disease progression and infection in *B. napus*, our RNA-seq analyses and histological examination of the infected tissues suggested that other cellular defense systems were also playing a role in the observed phenotype in BN1703.2 plants. BN1703.2 host defense pathways appear to play significant roles in host plant protection through quantitative upregulation of PR proteins, salicylic acid-mediated signalling and defense, fungal cell wall degrading enzymes and the production of secondary metabolites, compared to untransformed plants early in infection. Accumulation of defense-related mRNAs appears to be a common response within tolerant or resistant genotypes of *B. napus* in response to different pathogens [56]. For example, the *P. brassicae* resistant *B. napus* cultivar, Laurentian, activates salicylic acid-mediated defense responses, thaumatins, and PR proteins [56]. Additionally, the over-expression of *MITOGEN ACTIVATED PROTEIN KINASE 4 SUBSTRATE 1* in *B. napus* confers tolerance to *L. maculans* and accumulates transcripts associated with salicylic acid mediated defense [57]. Taken together, initial knockdown of a fungal mRNA facilitated by HIGS enhances innate defense of the host plant and provides additional levels of protection against pathogen infection over time.

Inducible anatomical barriers formed at or around the xylem of the host plant provide an additional level of protection that prevents the movement of the advancing fungus [58]. Traditionally, these barriers have been observed in plants infected by vascular wilt pathogens such as *Fusarium sp*. and *Verticillium sp.* [59–62]. Here, we observed both horizontal and vertical restrictions within cross-sections of *S. sclerotiorum* infected stems seven days post infection. However, horizontal restrictions through the formation of the vascular coating and deposition of callose are a structural barrier more likely to prevent radial mycelial movement towards the pith. Callose has been described as both a structural barrier against fungal pathogens and a cellular matrix that can accumulate antimicrobial compounds [63]. The deposition of callose in BN1703.2 like slows progression of *S. sclerotiorum* into medullary rays. Furthermore, the electron dense band running parallel to the site of infection likely serves as a second horizontal restriction coating cells of the secondary stem tissues. Vascular coatings occur at vessel walls, parenchyma cells and pit membranes to reinforce and create a dense, amorphous layer [58]. The composition of the coating varies, depending on the host and pathogen, but phenolics are the primary compounds, and phenolic polymers like lignin or suberin are the principal multi-molecular components [64]. Upregulation of the entire suberin biosynthesis pathway within BN1703.2 stems together with positively staining Sudan IV regions in the xylem, suggests that the cellular coating is at least partially suberized. Suberin is a complex polyester biopolymer consisting of polyphenolic and polyaliphatic domains, and it is not normally produced within the stem [65]. However, in many plant species, it has been shown that suberin production is induced during pathogen attack [66]. The deposition of suberin likely acts as a cellular obstruction to pathogen-derived metabolites such as pathogenicity factors and cell wall degrading enzymes, while still allowing plant-derived defense factors to be secreted [67]. A vascular coating was also observed within *B. napus* cv. ZY821, a moderately tolerant cultivar to *S. sclerotiorum*. Its tolerance is controlled through the activation of glucosinolate biosynthesis, which slows pathogen progression. It is possible that the formation of this vascular coating in *B. napus* in response to *S. sclerotiorum* infection is facilitated by the diminished pathogen advancement induced by the hpRNA or glucosinolate production.

Taken together, this study provides evidence that HIGS-based control of *S. sclerotiorum* in *B. napus* contributes to and complements innate cellular defense mechanisms of the host plant to reduce disease symptoms. Our results show that the reduction in disease symptoms was not solely the result of a hpRNA targeting *ABHYDROLASE-3* of *S. sclerotiorium*. Instead, activity of the hpRNA appears to provide *B. napus* with the necessary time required to activate innate global defense gene activity. While this approach was successful in protecting *B. napus* against *S. sclerotiorum*, a number of unanswered questions remain. For example, how is the transgene-derived RNA processed, sorted and taken up by the pathogen? Are long hpRNAs trafficked to the fungus in a similar mechanism as siRNAs and how can this dsRNA/siRNA transfer system be enhanced to confer additional levels of durability in economically important crops? Answering these questions along with others will ensure further development of this species-specific alternative to traditional fungal control and provide a path towards a more environmentally sustainable agriculture system.

## Materials and Methods

### *B. napus* growth conditions

Seeds of *B. napus* cv. Westar and the resultant transgenic lines were germinated in germination pouches (Phyto AB Inc., San Jose, CA, USA) and transplanted at the cotyledon stage into Sunshine Mix No. 1 (SunGro Horticulture, Agawam, MA, USA) at 22 °C and 50–70% humidity within greenhouse conditions. The plants were grown until the 30% flowering stage for use in infection experiments.

### Preparation of Agrobacterium tumefaciens

Primers for cloning SS1G_01703 can be found in Table S1. Cloning into *Agrobacterium tumefaciens* followed the Gateway cloning protocol (Invitrogen, Carlsbad, CA, US). Target gene sequences were amplified using Phusion Taq (Thermo Scientific, Waltham, MA, US) under the following conditions: 98°C for 30 s; 35 cycles of: 98°C for 10s, 57°C for 30s, and 72°C for 30s; and a final extension of 72 °C for 5 min. Amplicons were gel purified (New England Biolabs, Ipswich, MA, US) and digested using FastDigest KpnI and XhoI (Thermo Scientific, Waltham, MA, US) according to the manufacturer’s protocols. The products were ligated into pENTR4 vector (Invitrogen, Carlsbad, CA, US) using T4 ligase (Invitrogen, Carlsbad, CA, US) according to the manufacturer’s protocol. Prepared plasmids were used to transform *Escherichia coli* MachI cells (Thermo Scientific, Waltham, MA, US), selected using 50 mg/L chloroamphenicol (MP Biomedicals Inc., Santa Ana, CA, USA) + LB solid media (Difco Laboratories, Inc, Detroit, MI, USA), and sequence inserts were confirmed using Sanger sequencing (The Centre for Applied Genomics, Toronto, ON, Canada). Once sequence was confirmed, Gateway LR clonase II reactions (Invritogen, Carlsbad, CA, US) with an input ratio of 4:1 (pENTR4:pHellsgate8) were used to shuttle inserts into pHellsgate8 (Invitrogen, Carlsbad, CA, US). Prepared plasmids were used to transformed *E. coli* MachI cells (Thermo Scientific, Waltham, MA, US). Colonies were selected using 50 mg/L spectinomycin (MP Biomedicals Inc., Santa Ana, CA, USA). To confirm transformation, colony PCR screens were conducted using GoTaq® Green Master Mix (Promega, Madison WI, USA) with the following conditions: 94°C for 2 min.; 35 cycles of: 94°C for 30 s, 55°C for 30 s., 72°C for 35 s.; final extension of 72°C for 5 min. Plasmids were isolated from positive colonies using Monarch Miniprep kit (New England Biolabs, Ipswich, MA, US) and then FastDigest XbaI or XhoI separately (Thermo Scientific, Waltham, MA, US) according to the manufacturer’s protocols to confirm double recombination. Plasmids were transformed into *Agrobacterium tumefaciens* GV3201 (Thermo Scientific, Waltham, MA, US) and selected using 50 mg/L spectinomycin and 50 mg/L gentamycin to maintain the pMP90 plasmid (MP Biomedicals Inc., Santa Ana, CA, US). A colony PCR was performed as stated above with the initial 94°C step for 10 min to confirm transformation.

### *B. napus* transformations

Cotyledons were excised at the petiole of five-day old *B. napus* cv. Westar seedlings. The cut petioles were dipped into a solution of transformed *Agrobacterium* (O.D. of 0.7 at 600 nm). Generation of calli, leaves, shoots and roots on the explants was induced through media transfers as described in Bhalla and Singh [68]. After root regeneration, the explants were transplanted to soil and allowed to grow and mature to produce T1 seed.

### DNA extraction and genotyping

DNA was extracted from leaf samples of 7-day-old *B. napus* using a GenoGrinder 2000 (Spex CertiPrep, Metuchen, New Jersey, USA) and DNA extraction buffer [0.25M NaCl, , 1M 2-amino- 2-(hydroxymethyl)-1,3-propanediol (TRIS)-HCl pH 7.5, 25mM EDTA pH 8, 0.5% SDS]. DNA was precipitated with isopropanol, washed with 75% ethanol, and suspended in water.

PCR was performed using GoTaq® Green Master Mix (Promega, Madison WI, USA) according to the manufacturer’s instructions using SS1G_01703 specific primers (Table S1) . Thermocycler conditions were as follows: 94°C for 2 min.; 35 cycles of: 94°C for 1 min., 56°C for 30s., 72°C for 35s.; final extension of 72°C for 5 minutes. PCR products were loaded on a 1% agarose gel containing ethidium bromide. 2μL of sample was loaded per well with FastRuler Middle Range DNA Ladder (Thermo Fisher) as reference to compare band size. Gels were run for 18 minutes at 100 volts and 200 mAh. Gels were visualized under UV light using an Axygen® Gel Documentation System (Axygen, Corning, NY, US).

### Copy number determination

The number of transgene copies present within T1 explants was determined via quantitative RT PCR using BNHMG I/Y as the endogenous reference gene as described in Weng *et al.* [43] (Table S2). The cycle threshold values of the DNA of our transgene were compared to that of the endogenous reference gene through the formula:

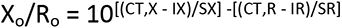

Where X is the transgene, R is the endogenous control, I is the intercept of a standard curve dilution series for each primer set and S is the slope of the standard curve.

### RNA isolations

RNA was extracted from infected and uninfected plant tissue. For infected leaves, a circular punch (30 mm diameter) was taken to collect the necrotic lesion and surrounding leaf tissue. For infected stems, cross-section disks (10 mm wide) were taken from the site of infection. The tissue was ground in an RNase-free mortar and RNA extracted using the Qiagen RNeasy Plant Mini kit (Toronto, ON, Canada). RNA was treated with Qiagen RNAse free DNAse (Toronto, ON, Canada), then cDNA synthesized using Quantabio qScript cDNA (Beverly, MA, USA). Quantity and purity were assessed spectrophotometrically and the quality of RNA samples verified using electropherogram profiles and RNA Integrity Numbers (RIN) (All samples had a RIN greater than 8.5) with an Agilent 2100 Bioanalyzer and RNA Pico Chips (Agilent, Santa Clara, CA, USA).

### RT qPCR

cDNA was diluted 1:40 in DNAse-free water for all qPCR reactions unless otherwise specified. *B. napus* and *S. sclerotiorum* primers were designed using Primer BLAST (www.ncbi.nlm.nih.gov/tools/primer-blast) to PCR amplify gene fragments ranging between 180 and 200 bp in length and Primer3 (http://bioinfo.ut.ee/primer3/) was used to evaluate potential primers [69]. A complete list of primers and their efficiencies can be found in Table S2. mRNA transcript abundance was determined using qPCR on the Bio-Rad CFX96 Connect Real-Time system using SsoFast EvaGreen Supermix (Bio-Rad Laboratories, Hercules, CA, US) in 10 μL reactions according to the manufacturer’s protocol under the following conditions: 95°C for 30s, and 45 cycles of: 95°C for 2s and 60°C for 5s. Melt curves with a range of 65 – 95°C with 0.5°C increments were used to assess nonspecific amplification, primer dimers, and aberrant amplifications. Relative accumulation was calculated using the ΔΔCq method, relative to Sac7 (SS1G_12350) for *S. sclerotiorum* and BNATGP4 (*BnaC08g11930D*) for *B. napus*.

### Linear-hairpin variable primer RT-qPCR

Linear-hairpin primers were designed to target select small RNAs as described by Lan *et al.* [70]. SsoFast EvaGreen Supermix (Bio-Rad Laboratories, Hercules, CA, US) was used for qRT PCR with conditions of 94 °C for 4 min, followed by 50 cycles of 94 °C for 30 s and 56 °C for 30 s. Primer sequences can be found in Tables S1 and S2.

### Leaf infection assays

*S sclerotiorum* ascospores were collected at the Morden Research and Development Centre, Agriculture and Agri-Food Canada, Morden, MB, Canada and stored at 4°C in desiccant in the dark. Plants were infected when they reached the 30% flowering stage. *S. sclerotiorum* ascospores (1 x10^6^ mL^-1^) were suspended in a 0.02% Tween80 (Sigma-Aldrich, St. Louis, MO, USA) solution. 10 µL of the ascospore solution was transferred onto senescing *B. napus* petals in a petri plate and sealed with Parafilm. Ascsospore-inoculated petals were stored at room temperature (21°C) for 72 hours and allowed to germinate prior to being inoculated on the leaf surface. The infected petal was placed onto the surface of a host leaf and sealed in a zip-lock bag to maintain humid conditions. After two days, lesion sizes were quantified using ImageJ software (imagej.nih.gov).

### Stem inoculations

A 3 x 3 mm cube of actively growing *S. sclerotiorum* from the leading edge of hyphae on a potato dextrose agar plate was collected using a syringe tip and placed onto the stem of a mature *B. napus* plant 30-50 mm above the soil. The fungus was sealed to the stem using a 30 x 20 mm strip of parafilm and lesion length was measured seven days post inoculation.

### Whole plant spray inoculations

Plants were infected when they reached the 30% flowering stage. *S. sclerotiorum* ascospores (1 x10^6^ mL^-1^) were suspended in a 0.02% Tween80 (Sigma-Aldrich, St. Louis, MO, USA) solution. 25 mL of spore solution was sprayed onto the whole surface of each plant using a spray bottle. A 4x6 ft humidity chamber was constructed and covered with vinyl plastic sheeting. Once sprayed, the plants were placed within the humidity chamber and the chamber was heavily misted to maintain high humidity. The plants remained in the chamber for a seven-day period, at which point they were removed and the infection was allowed to continue to completion on the greenhouse benchtop.

### mRNA library synthesis, sequencing and bioinformatic analysis

RNA was extracted as described above and cDNA libraries were constructed by Genome Québec according to their mRNA stranded library protocol. The Illumina HISeq4000 was used to generate 100 bp PE reads. FastQ files were trimmed using Trimmomatic 0.33 [71]: adapter sequences, initial 12 bases of raw reads, low quality reads with a quality score under 20 over a sliding window of 4 bases and reads with an average quality score under 30 removed during the trimming process. Remaining reads shorter than 40 nucleotides were also removed. The high sensitivity mapping program HISAT2 [72] was then used to align trimmed reads to the *B. napus* and *S. sclerotiorum* genomes [47, 48]. The mapped reads were sorted with Samtools [73] and assigned to genes with featureCounts [74]. EdgeR [75] was used to perform differential gene expression analysis on raw counts using a log2 fold change and FDR cutoff of |2| and <0.01, respectively. All RNA-seq data exploration were performed in R studio. The Principal Component Analysis was carried out using plotPCA function in DESEq2 [76] package. Stacked bar and Upset plots were generated in R [77]. GO term enrichment was carried out using SeqEnrich to identify enriched processes and the genes associated with them [49].

All data have been deposited at the Gene Expression Omnibus (GEO): GSE 184812.

### Small RNA library synthesis, sequencing and analysis

Total RNA was extracted as described above and cDNA libraries were constructed by Genome Quebec according to their small RNA library protocol. The Illumina HISeq4000 was used to generate 100 bp PE reads. FastQ files were trimmed using Trimmomatic 0.36 [71]: adapter sequences, initial 12 bases of raw reads, low quality reads with a quality score under 20 over a sliding window of 4 bases, and reads with an average quality score under 30 or under 18 bp removed during the trimming process. Bowtie was then used to align the SS1G_01703 transgene to the small RNA dataset [78].

### Light microscopy

Tissues were embedded in historesin (Leica, Wetzlar, GER). Briefly, tissue was vacuum infiltrated in 2.5% glutaraldehyde/1.6% paraformaldehyde fixative. Methylcellosolve (Sigma-Aldrich, St. Louis, MO, USA) was used to remove pigment and three transfers of 100% ethanol every 24 hrs for three days was used to completely decolourize and dehydrate the tissue.

Historesin was then infiltrated into the tissue during a three-day period of increasing concentrations of the resin. Day one was 1:3 activated historesin:100% alcohol, day 2 was 2:3 activated historesin:100% alcohol and day 3 was 100% historesin. The tissue was then embedded into blocks with an embedding solution of activated historesin (Leica, Wetzlar, GER), hardener (barbituric acid) and polyethylene glycol. Sections were cut at 3 μM and stained with lactophenol ‘cotton’ blue (to detect fungal chitin) and counterstained with 0.1% Safranin O for 20 minutes and 5 seconds respectively, aniline blue (to detect callose) and Periodic-Acid Schiff’s (to detect polysaccharides) for 10 minutes and 15 minutes for both Periodic-Acid and Schiff’s base respectively or 0.1% toluidine blue O (to detect lignin) and Periodic-Acid Schiff’s for 15 minutes and 15 minutes for both Periodic-Acid and Schiff’s base respectively. Hand sections of tissue were also cut and stained with Sudan IV (Sigma-Aldrich, St. Louis, MO, USA) (to detect suberin) for 20 minutes and cleared in deionized water. Slides were imaged using a Leica DFC450C camera. Three replicates (stems from individual plants) were viewed of both RNAi and untransformed lines and a representative image was published.

### Scanning electron microscopy

Stem cross-sections from 7-day old infections were collected and imaged immediately on the Hitachi TM1000 tabletop system using environmental SEM. Images were taken quickly after sealing the chamber, as to not allow the samples to dry out. Three replicates (stems from individual plants) were viewed of both RNAi line and untransformed and a representative image was published.

## Acknowledgments

We would like to thank Austein McLoughlin for his help with the cloning of the pHELLSGATE8 vector, Dr. Ravinder Sidhu for her help with the microscopy, and Nina Huynh for her help with plant transformations. This work was generously supported by grants from the province of Manitoba Agricultural Rural Development Initiative, the Canola Council of Canada, and the Western Grains Research Foundation to S.W. and M.F.B., N.W. was supported by a National Science and Engineering Research Council Graduate Scholarship and the Manitoba government Tri Council Top-Up award in addition to the University of Manitoba Graduate Fellowship.

## Contributions

N.W. performed bioinformatic analyses, primer design, RNA extractions, qRT PCR, plant transformation, *in planta* assays, and manuscript preparation. D.J.Z. performed bioinformatic analyses and microscopy. P.L.W. performed bioinformatic analyses. D.S.S. and K.T.B. performed *in planta* assays. D.K. performed plant transformation. S.K.S and O.W. performed bioinformatic analyses and manuscript preparation. S.W. and M.F.B. conceived the ideas and prepared the manuscript.

## Supporting Information

**Fig. S1 Agarose gel images of the 1703 transgene fragment in BN1703 lines.** Positive control uses the 1703:pHELLSGATE8 construct.

**Fig. S2 Small RNA sequence reads aligning to the 1703 transgene**. Reads were plotted to their respective location upon the transgene sequence using Geneious Prime software (https://www.geneious.com) to identify highly abundant molecules.

**Fig. S3 Relative leaf and stem lesion sizes and PR1 transcript abundance of BN1703.2 infections compared to untransformed.** (a) Asterisks represent statistical differences from the untransformed control (one-tailed t-test with Bonferroni correction, p<0.05). (b) Samples were normalized to the housekeeping reference BNATGP4. Data represents 3 biological replicates with error bars representing standard error. Asterisks represent statistical differences from the untransformed control (one-tailed t-test with Bonferroni correction, p<0.05).

Fig. S4 PCA clustering analysis of *B. napus* (A) and *S. sclerotiorum* (B) aligned reads of infected untransformed and BN1703.2 stems 0, 1 and 7 days post-inoculation.

**Fig. S5 Heatmap of a subset of differentially expressed genes in untransformed and transgenic BN1703.2 infected stems 7 days post-inoculation compared to 1 dpi post infection.** Genes were involved in *S. sclerotiorum* infection common to the biotrophic and necrotrophic stages of infection (false discovery rate<0.05).

**Fig. S6 Scanning electron and light microscopy stem cross-sections stained with toluidine blue, Sudan IV, and aniline blue comparing transformed and untransformed plants** (A) Scanning electron micrograph of a stem cross-section of the semi-resistant cultivar of *Brassica napus* cv. ZhongYou 821 7 days post-inoculation. Black arrowheads indicate *S. sclerotiorum* hyphae and white indicates the vascular coating. Scale bar represents 0.5 mm. (B) Toludine- blue and Periodic-Acid Schiff’s staining of BN1703.2 infected stem cross-sections 7 days post inoculation. Black arrowheads indicate negative detection to the vascular coating deposits within the secondary xylem by these stains. Scale bar represents 50 μm. Scanning electron micrographs of uninfected untransformed (C) and BN1703.2 (D) stem cross-sections. Scale bars represent 0.5 mm. Hand-cut stem transverse sections stained with Sudan IV of uninfected untransformed (E) and BN1703.2 (F) stems. Uninfected stems exhibit low affinity for Sudan IV in both. Scale bars represent 250 μm. Uninfected untransformed (G) and BN1703.2 (H) stem cross-sections stained with aniline blue and periodic acid-Schiff’s reagent. Aniline blue staining is scant in both cultivars, indicating low callose deposition. Scale bars represent 250 μm.

Data S1 Summary of the trimming and alignment statistics for all samples from the RNA sequencing analysis, the counts data for both *B. napus and S. sclerotiorum* gene alignments, enriched GO terms from SeqEnrich analysis and the total small RNA sequences aligning to the 1703 transgene found through small RNA sequencing.

